# Dynamic Measurements of Bovine Serum Albumin Aggregation Using Small Angle Neutron Scattering

**DOI:** 10.1101/368092

**Authors:** James I Austerberry, Daniel J Belton

## Abstract

The rapid and complex nature of protein aggregation makes the identification of aggregation mechanisms and their precursors challenging. Here we demonstrate the novel use of small-angle neutron scattering to perform dynamic real-time measurement and analysis of protein aggregation. Changes in bovine serum albumin monomer population and aggregate size are identified at several isothermal temperatures. Kratky plots indicate that the aggregation of BSA occurs through the partial unfolding of the monomer. Dual population modelling of the scattering data indicates that the protein nucleates and grows through a two stage mechanism; a rapid burst phase and a slower growth phase. Both stages are observed to follow Arrhenius behaviour between 70-80 °C.

## Introduction

Protein aggregation is the self-association of protein monomers to form higher order structures. Understanding protein aggregation is important for many areas, including therapeutic protein processing and formulation, elucidating the mechanism of neurodegenerative diseases, and developing novel biomaterials. For example, preventing aggregation of therapeutic proteins is essential, since it can lead to a loss of product, reduce efficacy, alter biological activity and pharmacokinetics, and even raise safety concerns such increased immunogenicity [1–4].

Protein aggregation can be induced by various processing steps and solution conditions, such as over-expression, refolding, chromatography, freeze-thaw, agitation, foaming, temperature, pH, salinity and additives [5–7]. The resulting aggregates can take on a number of different structures depending on the solution conditions. Well-established structures include particulates [8], fibrils [9] and spherulites [10]. Solution pH is considered to be of particular importance in controlling aggregate morphology, with pH values close to the isoelectric point favouring the formation of particulates [8].

The literature is awash with studies that postulate different empirical and mechanistic models of protein aggregation [11–14]. Mechanistic models have been broadly categorised as monomer addition, reversible association, prion aggregation, minimalistic 2-step or quantitative structure-activity relationships. In addition, *Roberts et al* proposed a comprehensive generic mechanistic scheme for protein aggregation with associated generic equations, which can be simplified for certain conditions and limiting cases [15]. Overall, most models are based on the concept of aggregation via a reactive/denatured monomer which is in equilibrium with a native state; the reactive/denatured monomer is able to aggregate, initially via a nucleation step followed by a growth phase.

The temperature dependence of these processes is not straight forward and it is generally recognised to be non-Arrhenius [5, 16]. The various stages involved in aggregation may be affected by temperature in different ways and to different degrees. For example, collision dependant processes will be affected by changes in diffusion and collision frequency, whereas temperature dependant unfolding processes will be affected by changes in hydrophobic interactions and the stability of weak interactions holding the polypeptide chain fold together. The thermodynamics of unfolding itself is also complicated but a generalised relationship between the Gibbs free energy of protein unfolding and temperature has previously been described [17, 18] [19].

A wide range of techniques have been used to study protein aggregation, including turbidity, static light scattering, dynamic light scattering, fluorescence spectroscopy, light scattering spectroscopy, circular dichroism, infrared spectroscopy, size-exclusion chromatography, analytical ultracentrifugation, gel electrophoresis, capillary electrophoresis, mass spectrometry, calorimetry, electron microscopy and small-angle X-ray scattering; several reviews and summaries are available [20–22]. Measurements can be either in-situ or ex-situ. For example, methods requiring sample preparation such as separating, drying, fixing or labelling, are necessarily ex-situ and limited to static measurements, whereas in-situ methods offer the possibility of real-time analysis. Furthermore, mechanistic studies necessitate that measurements be related back to a real change in aggregation state, such as aggregate size or monomer depletion, and studies using ex-situ approaches require the aggregation process to be reliably halted and persevered for a series of samples representing different time points.

The growth phase of aggregation would appear relatively easy to monitor and analyse, requiring only the analysis of particle growth rate [23, 24] and often reducing to simple first order behaviour [15]. Whereas, monitoring the nucleation stage of aggregation remains more difficult, due in part to the difficulty of isolating nuclei for analysis and limitations of techniques available for direct characterisation of nuclei [25]. Furthermore, the structure of the nucleus is not clearly defined in the literature [5]. Overall, it would appear that the majority of studies are focused on aggregate growth, with nucleation processes seemingly elusive and limited only to inference [25] or discussion of lag phases and induction times [5, 26, 27].

Small-angle neutron scattering (SANS) has been successfully used to analyse static protein aggregate samples [28–30], but has been somewhat overlooked by general reviews of techniques for analysing protein aggregation. Particular advantages include probing structures over a range of length scales and minimal beam damage to samples. Here we present the use SANS to examine the formation and growth of particulate aggregates of bovine serum albumin (BSA) using real-time in-situ measurements and derive information on the aggregation prone state and aggregate growth mechanism.

## Results and Discussion

In order to examine sample quality and ensure the absence of any aggregates in the initial solution, a scattering profile for 2 mg/ml BSA in 90 % D_2_O pD 7.8 was obtained at 25 °C before aggregation was initiated (Fig 1). A fit for BSA was applied to the scattering data using CRYSON; a program for evaluating small-angle solution scattering from macromolecules with known atomic structure [31, 32]. This is based on crystal structure 3V03 in the protein data bank [32]. This yielded an excellent fit, indicating that the sample was initially monomeric and without aggregate contaminant prior to the heat treatment. The CRYSON data fit reveals an 80% deuteration of available exchange atoms, where the remaining protons are not accessible by the solvent. This is in agreement with previous studies where a similar level of deuteration was observed [33]. The fit yielded values for the volume of BSA as 82666 Å3 and a radius of gyration of the envelope of 32 Å which agrees with literature vales [34].

**Figure 1.**
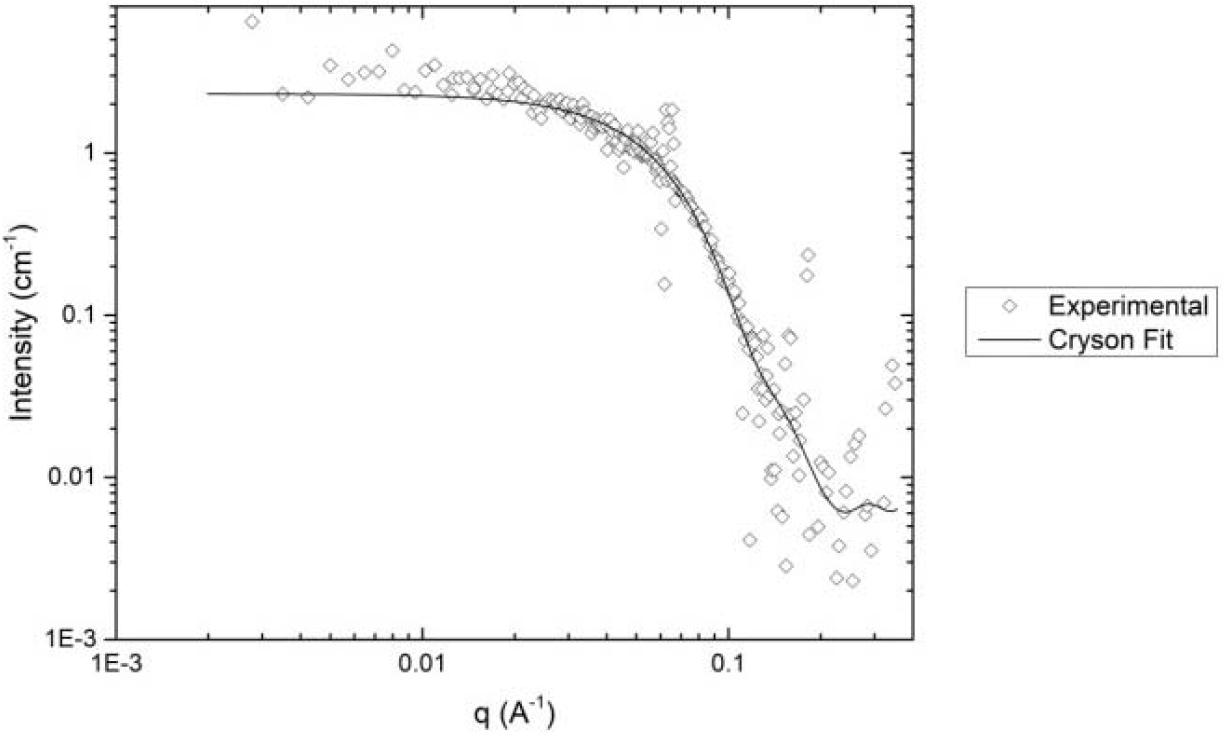
The scattering profile for 2 mg/ml BSA in 90% D2O pD 7.8 at 25 °C prior to thermally induced aggregation. The line shows the CRYSON fit to BSA scattering data based on crystal structure (Svergun 1995 and 1998).

Samples of 2 mg/ml BSA in 90 % D_2_O pD 7.8 were incubated isothermally at 70, 75 and 80 °C, with data collected continuously in one minute accumulations. Figure 2A shows the changes in SANS profile at selected intervals during the aggregation process at 80 °C. At t=0; in the native state the intensity at low q values is low (15 cm^-1^). Increasing incubation time results in increasing intensity (>100 cm^-1^), at low q indicating the formation and growth of large scale aggregates over time. The fitting of Eq.1 to the scattering plot *I(q)* yields an *I0* value which is directly related to the y-intercept: Equation 1

**Figure 2A.**
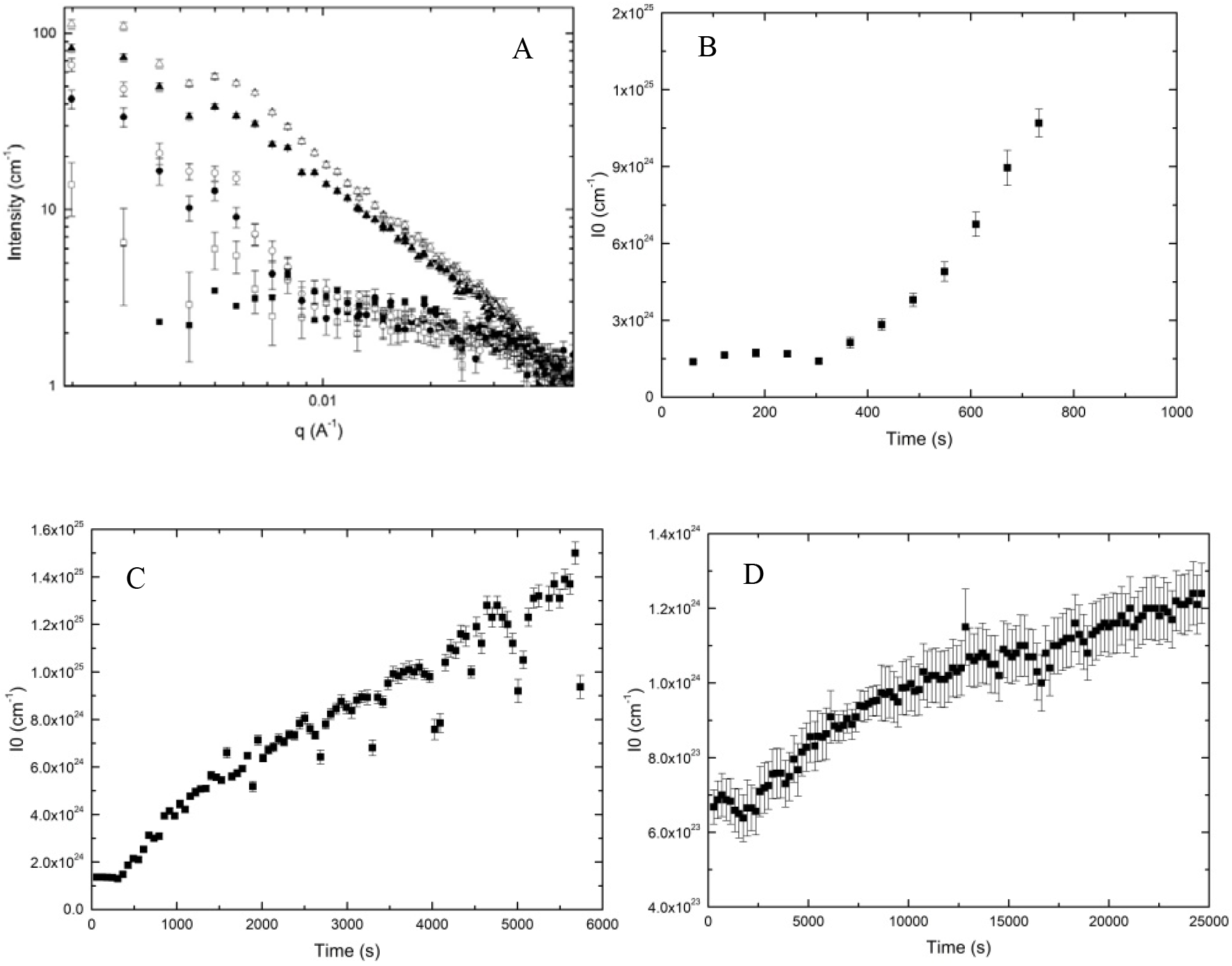
Change in BSA scattering data (I vs. q) over time at 80°C. 60 s (closed squares), 600 s (open squares), 1200 s (closed circles), 1800 s (open circles) 2400 s (closed triangles), 3000 s (open triangles). 2B-D shows the scaled I0 values for 80 °C (B), 75 °C (C), and 70 °C (D) respectively. Note the different time scales on the x-axis.

Where Δρ is the difference in scattering length density between protein and buffer, *v* is the volume of the protein in solution. Figures 2B-D illustrate the I0 values at each time interval for the 80, 75 and 70 °C experiments respectively. At 80 °C the rate of increase in I0 occurs more rapidly (300 s) than at 75 °C (500 s) or 70 °C (2500 s). As would be expected this indicates that the higher temperatures are increasing the rate at which BSA aggregates are forming. The increase in aggregation could be explained by the increasing the population of unfolded states of the BSA monomer combined with the increased kinetic energy resulting from the elevated temperatures [35].

In order to assess the level of folding present during aggregation, a temperature scan for BSA was obtained between 25 and 75 °C. Figure 3 shows a Kratky plot for the 2 mg/ml BSA in 90 % D_2_O pD 7.8 sample at select increasing temperatures. The presence of a defined peak in the Kratky plot at high q where the plot tends towards q^-4^ (q > 0.1 Å^-1^) is evidence of a folded protein, as is seen at 25 °C, whereas an unfolded chain would exhibit no defined peak in the q^2^ I value as the high q values tend toward q^-2^ [36]. As temperature is increased, the q^2^ I value at high q increases. This is indicative of the structure of the protein becoming unfolded, however the presence of the peak at 75 °C indicates that only partial unfolding has occurred [37]. This indicates that partial structural disruption of the monomer is necessary to initiate aggregation, with the protein remaining largely globular in this aggregation prone state and subsequent aggregate. This correlates well with the literature which indicates that complete unfolding of BSA may not occur until almost 90°C [38] whilst we observe that aggregation of the protein occurs at lower temperatures (Fig 2). We have observed the effect of partial unfolding initiating aggregation is not isolated to BSA [39, 40].

**Figure 3.**
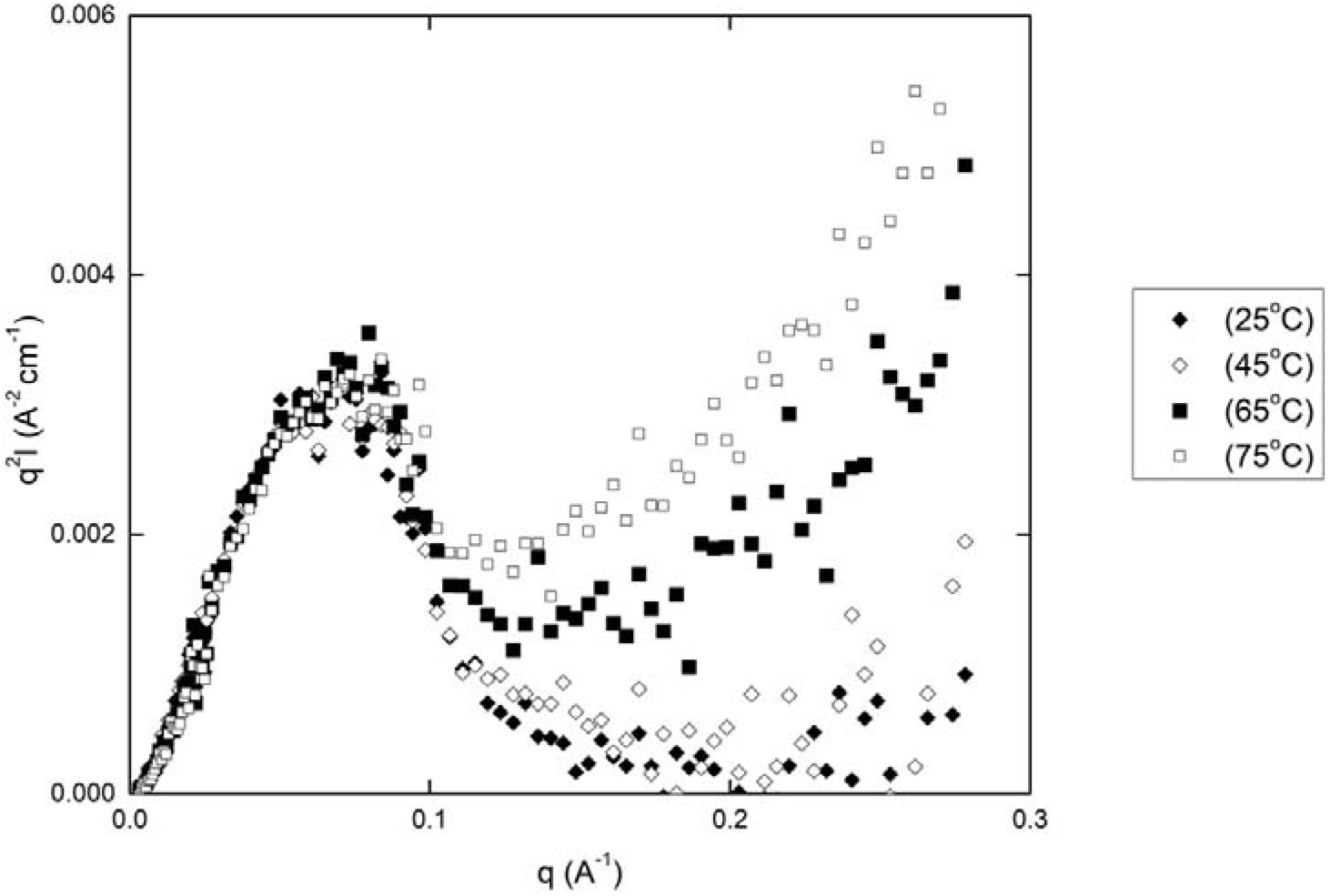
Kratky plot (q v q^2^I) of 2mg/ml BSA in 90 % D_2_O pD 7.8 at increasing temperature: 25°C (closed diamonds), 45°C (open diamonds), 65°C(closed squares), 75°C (open squares).

In order to assess the aggregation within the sample, a dual population model was fitted to the scattering data (Fig 4). The model is based on two populations of spherical particles, one representing the monomer population and the other the aggregates (see methods). The value for the radius of the monomer fraction was determined from fits of a single Rayleigh scatterer to scattering profiles of the BSA at t=0 (Supplementary Fig 1). This produced a radius of the monomer of 34.4 Å, which was included as a constrained value in the dual population fit. Whilst this does not take into account changes to the monomer during the heating process, we find that these changes in size are small when compared with the change in size resulting from aggregation of the BSA monomers. The model shows a good fit (R = 0.95) over the entire data range and allows the aggregate size and level of monomer present to be assessed. This assessment is based on two parameters within the model; monomer volume fraction, F_M_, and aggregate radius, R_A_. It should be noted that the dual population model fit is less reliable when the level of scattering contributed from the aggregate population is very low. Attempts to fit scattering from BSA prior to heating, where fitting with CRYSON demonstrates the protein is monomeric, does not yield the expected monomer fraction and the fitting parameter errors are considerably higher than for aggregated samples.

**Figure 4.**
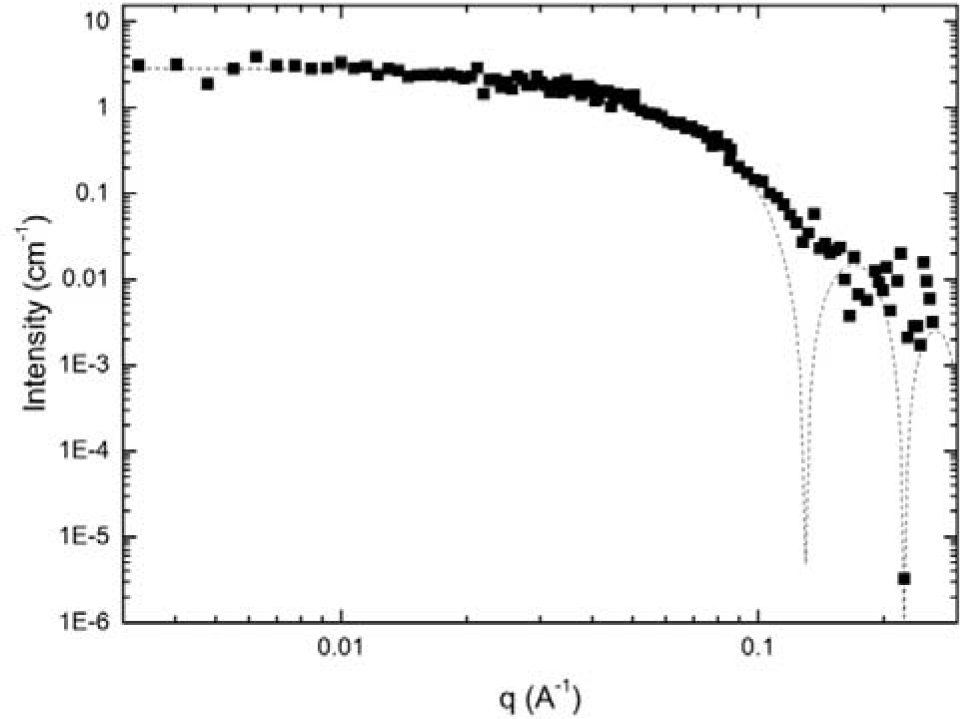
Intensity versus scattering vector for BSA after 20 minutes incubation at 75°C in 90% D_2_O pD 7.8. Dashed line represents the fit of the dual population model to the data.

Comparison of the growth of aggregate radius over time indicates that BSA undergoes a 2-step aggregation growth mechanism at this temperature range (Fig 5). The initial phase is represented by a rapid growth burst phase. At 80 °C the burst phase is identified as an increase of aggregate size to 700 nm after 1000 s, whilst at 75 and 70 °C the growth of 180 and 100 nm over 500 and 250 s respectively. This is followed by a significantly slower phase of growth than the first: at 80 °C the aggregate radius increases 20 nm over 500 s and 20 over 4000 s at 75 °C, whilst at 70 °C the growth is a near plateau. At each temperature, the aggregation kinetics exhibits a two stage aggregation process, which has been exhibited previously at similar temperature ranges [41, 42]. Extrapolation of the initial growth rates each plots indicates the initial detected aggregate radius to be ∼65 nm for each temperature. As this is temperature independent this value is a limit of the model and the ability of the fit to discriminate between scattering contributions from the monomers and aggregate. It may be possible to achieve a smaller initial aggregate size from scattering plots of increased signal to noise. The presence of two separate aggregate growth rates may be due to the change in secondary structure during the aggregation process. BSA is known undergo a decrease in α–helical content and an increase in β–sheet [41]. As β-sheet content is known in the formation of amyloid fibrils, where conformational restrictions and charge distributions limit the number of possible orientations monomer addition may occur, aggregation rates are known to be slower when fibril formation is occurring [43, 44]. This correlates with the slower rate of growth seen in the latter stages of each temperature run (Fig 5). Fig 5 also illustrates the temperature dependence of both aggregate growth phases on BSA, whilst the T_m_ of BSA has been shown to be around 90 °C (dependent of solution conditions) [38], whilst kinetics are also dependent on solution constituents [45]. At these temperatures the change in unfolded quantity of the monomer should be minimal, therefore the dependence on temperature is likely linked to minor unfolding events or the increase in kinetics of the system.

**Figure 5:**
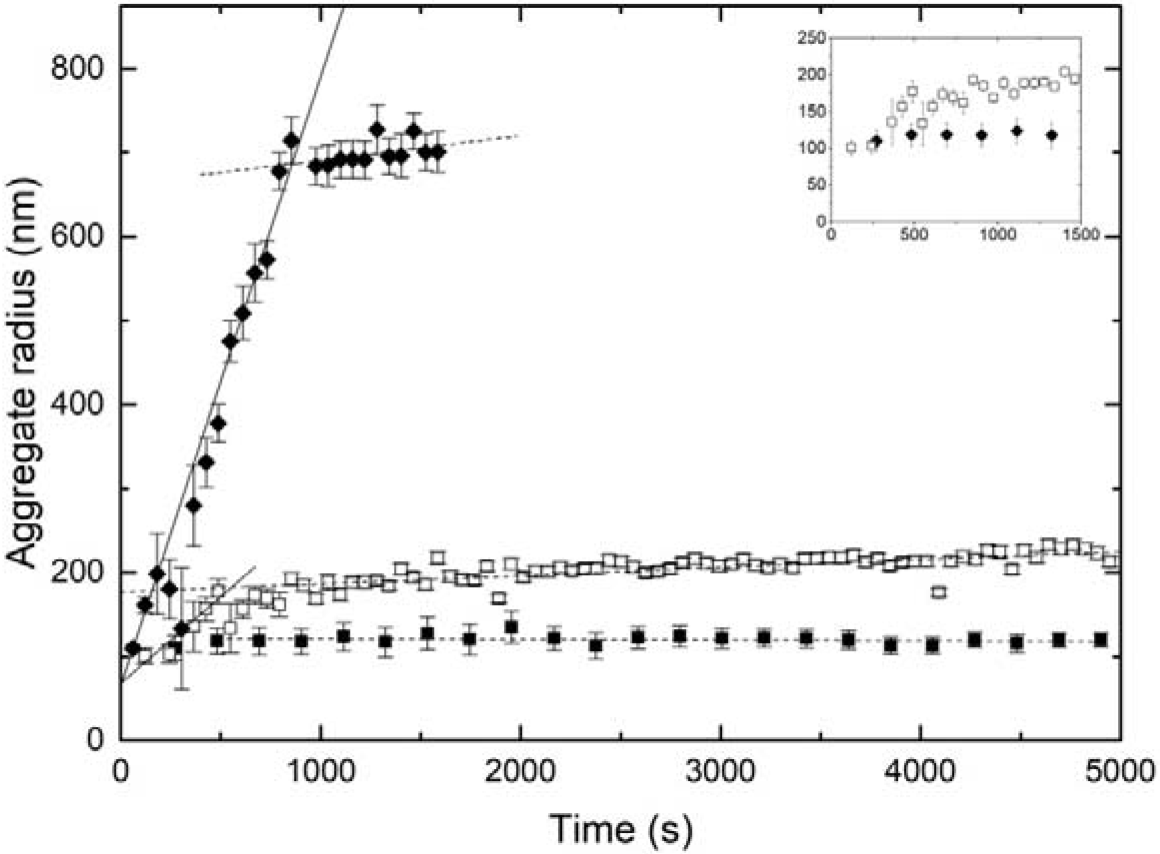
Aggregate radius over time from dual population model at 70, 75 and 80 °C in 90% D_2_O pD 7.8. Inset shows early burst phase for 70 and 75 °C.Linear fits are included to the burst phase (solid line) and growth phase (dash line).

Arrhenius plots were created for both the rapid burst phase and the slow growth phase of BSA aggregates using their respective growth rates (Fig 6). The differences in slope between the two plots further illustrate the large difference in rates between the two phases. The linear nature of the plots show that both phases exhibit Arrhenius behaviour (R >0.95). This indicates that both growth phases follow a first (or pseudo first) order reaction scheme. This is consistent with the emergence of a constant concentration of aggregate nuclei, and monomer addition which would yield a first order reaction rate as the rate is dependent on monomer concentration for both of the phases [46]. Linear fits of to both of the slopes, where growth rate, k = A*exp(-Ea/R*T) in plots A and B lead to pre-exponential factor being calculated as 5.06 and 5.15 s^-1^ respectively, whilst the activation energy for each is 111 kcal mol^-1^ and 254 kcalmol^-1^. The linear nature of the plots indicates that over this temperature range there is minimal change in the conformation of the BSA monomer. However this plot cannot be used to estimate the stability of the protein monomer at room temperature, as Arrhenius plots of protein aggregation are frequently non-linear over large temperature ranges [16]. These values are larger than those seen for fibrillar aggregate formation [47, 48] and comparable with those of globular protein aggregation [49–51] and indicate the relative temperature dependence of the BSA monomer against aggregation. The sizeable difference between the two stages of aggregation indicate a significant difference in the aggregation mechanism and a multi stage aggregation mechanism [16].

**Figure 6:**
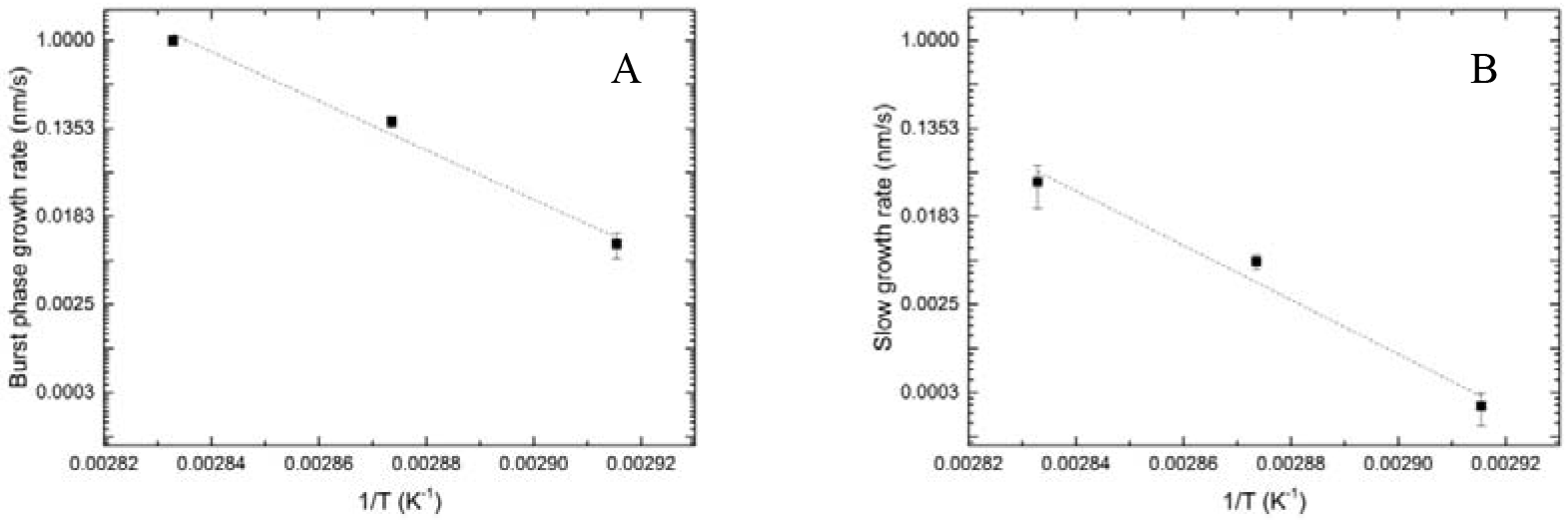
Arrhenius plots for burst phase growth (A) and slow growth rate (B) for 2 mg/ml BSA in 90 % D_2_O pD 7.8 at temperatures 70, 75 and 80 °C.

## Conclusions

Here we have demonstrated the use of small angle neutrons scattering as a tool for elucidating the mechanisms of aggregation in bovine serum albumin. Whilst basic scattering plots can be utilised to assess the level of folding present prior to aggregation, more complex modelling of the morphology and populations of both monomer and aggregate are required to extract mechanistic information. The spherical model used here represents a good fit for BSA due to its globular nature, but it remains to be seen whether more complex molecules such as antibodies can be fit in this way, or whether more sophisticated models are required. Similarly, this model relies on a dual population fit being suitable to describe the aggregation of the protein studied. Whilst this is the case here, it must be considered on a case by case basis. Nevertheless, this analysis shows promise for the progression of protein aggregation kinetics.

## Materials and Methods

### Samples

Bovine serum albumin was purchased from Sigma-Aldrich (France). For SANS measurements, the powder was dissolved in H_2_O to create a concentrated stock solution of the protein, before a 1:9 dilution of the stock into 100 mM Tris, 100 mM NaCl buffer made up with D_2_O to a final concentration of 2 mg/ml. The solution was adjusted to pD 7.8 using diluted HCl solution.

### Measurements

SANS spectra were recorded at the D22 instrument at the Instiuit Laue-Langevin, France. The protein solutions were filled into quartz cuvettes and individually heated to temperatures of 70, 75 and 80 ºC. Data were measured at three sample-detector distances (2, 4 and 11.2 m) at a neutron wavelength of 6 Å to cover a q range between 0.002 and 0.350 Å^-1^. Measurements were completed every minute for a period of between 1–8 hours.

### Data Analysis

Data reduction was performed using the program GRASP [52], including normalisation for sample transmission, and background subtractions using a quartz cuvette containing 90% D_2_O buffer as described above.

CRYSON [31] was used to fit the scattering data of the monomer to the crystal structure of the BSA molecule. This programme takes the data input in the form of the PDS file and creates a spherically averaged scattering pattern which takes into account the hydration shell and extent of deuteration present in the sample. CRYSON attempts to minimise the difference in the chi-squared value between the predicted and experimental data to give values for the hydration levels and envelope size of the protein.

Scattering from both monomer; *F_M_(q,)* and aggregate samples; *F_A_(q)*, were modelled as a Rayleigh spheres, leading to a dual population of spheres fit using an equation developed from these known principles. This is presented in equation 1: 
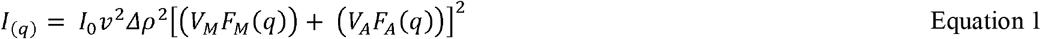
Where Δρ is the difference in scattering length density between protein and buffer, *v* is the volume of the protein in solution, *V_M_* and *V_A_* are the scattering volumes of the monomer and aggregates respectively and *I_0_* is a scaling factor. *F_M_(q)* and *F_A_(q)* are defined from the Rayleigh scattering resulting from the population of monomers and aggregates respectively: 
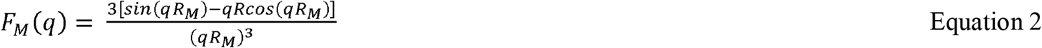

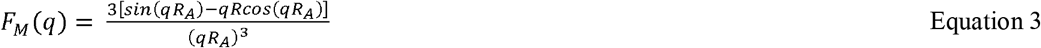

Where *R_M_* and *R_A_* are the radii of the monomer and aggregates.

## Supplementary

**Supplementary Figure 1:**
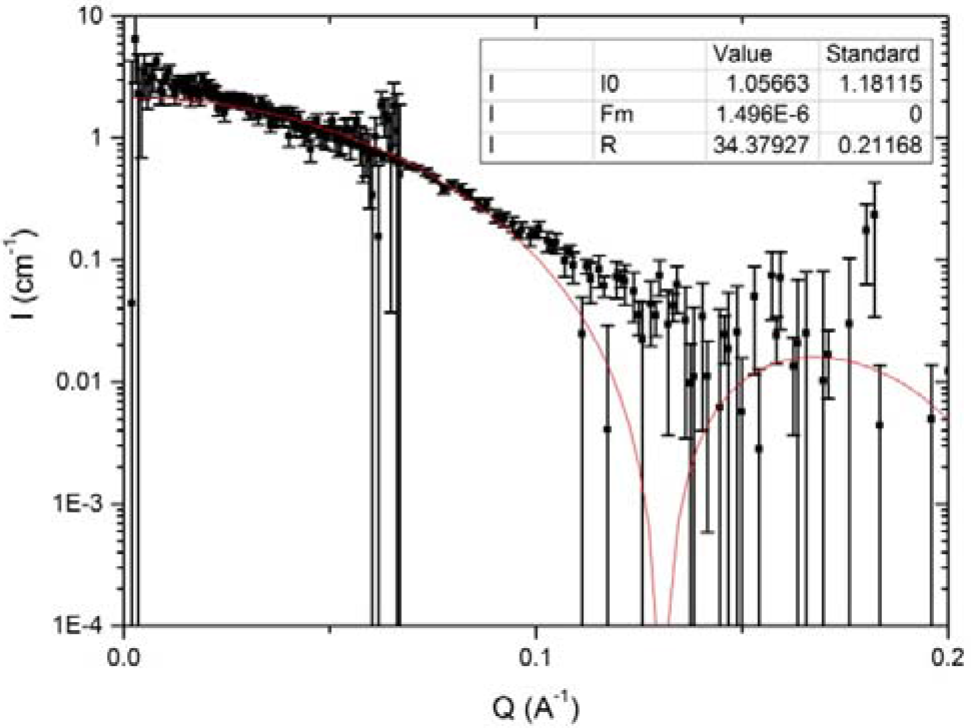
Fit of Eq 1 to 2 mg/ml BSA in 90% D2O pD 7.8 at 25 °C prior to thermally induced aggregation.

## Acknowledgements

The authors would like to thank Dr Peter Laity and Professor Susan Kilcoyne for valuable discussions, and the University of Huddersfield for funding.

The authors declare no conflict of interest within this work.

